# Assessment of urinary pharmacodynamic profiles of faropenem against extended-spectrum β-lactamase-producing *Escherichia coli* with canine *ex vivo* modeling

**DOI:** 10.1101/321836

**Authors:** Kazuki Harada, Takae Shimizu, Naoki Miyashita, Yoshiaki Hikasa

## Abstract

Extended-spectrum β-lactamase (ESBL)-producing bacteria are of great concern in companion animals with urinary tract infections (UTIs). Because of its high safety and stability in the presence of ESBLs, faropenem is assumed to be a candidate antimicrobial agent for canine UTIs with ESBL-producing bacteria. This study was performed to investigate the urinary pharmacokinetics and pharmacodynamics of faropenem administered at 5 mg/kg body weight in six healthy dogs using an *ex vivo* model. Six UTI pathogenic strains of ESBL-producing *Escherichia coli* (ESBL-EC) with the following faropenem minimum inhibitory concentrations (MICs) were used: 1 µg/mL (n = 2), 2 µg/mL (n = 2), 4 µg/mL (n = 1), and 16 µg/mL (n = 1). Urine samples were obtained every 4 h for the first 12 h after faropenem administration for measurement of the urine drug concentration and urinary bactericidal titers (UBTs). The urine concentration of faropenem peaked at 0 to 4 h after administration, with a mean maximum concentration of 584 μg/mL, and markedly decreased at 8 to 12 h (23 μg/mL). The median UBTs for all tested ESBL-EC strains were highest at 0 to 4 h and then significantly decreased at 8 to 12 h. These findings indicate that administration of faropenem more than once daily is recommended for the treatment of ESBL-EC-related UTIs in dogs. In addition, the median areas under the UBT–time curves (AUBTs) were significantly inversely correlated with the corresponding MICs for faropenem in the tested strains (*P* < 0.05). Notably, the median AUBTs were significantly higher in ESBL-EC strains with an MIC of 1 µg/mL than in those with an MIC of ≥4 µg/mL (*P* < 0.05). The present study serves as the basis of clinical application of faropenem for ESBL-producing bacteria-related UTIs in dogs.

## Introduction

Urinary tract infections (UTIs) are common bacterial infections in dogs, occurring in approximately 14% of dogs in their lifetimes with a variable age at onset [1]. *Escherichia coli* is the most common infectious bacteria, although various gram-negative and gram-positive bacteria can cause UTIs in dogs [2,3].

The prevalence of extended-spectrum β-lactamase (ESBL)-producing bacteria is of great concern worldwide in companion animals with UTIs [4,5]. Although ESBLs are usually involved in resistance to oxyimino-cephalosporins, penicillins, and narrow-spectrum cephalosporins, ESBL-producing bacteria are often resistant to other classes of antimicrobials [6]. These multidrug-resistant phenotypes of ESBL-producing bacteria have major implications in the selection of adequate empirical therapy regimens [6].

We previously reported high *in vitro* efficacy of several antimicrobials against ESBL-producing *E. coli* (ESBL-EC) isolates from companion animals [7]. Of these antimicrobials, faropenem is representative of the penem class and exhibits stability to hydrolysis by ESBLs [8,9]; additionally, it is highly safe in dogs [10,11]. These findings indicate that faropenem may be a promising candidate antimicrobial for canine UTIs with ESBL-producing bacteria. However, the urinary pharmacokinetic/pharmacodynamic profile, which is essential for assessment of treatment efficacy of antimicrobials in UTIs [12–14], remains to be investigated.

In the present study, we used liquid chromatography–mass spectrometry (LC-MS) to investigate the urinary pharmacokinetics of faropenem in dogs. We also measured urinary bactericidal titers (UBTs) and related parameters of faropenem against ESBL-EC strains from canine UTIs.

## Materials and Methods

### Sampling of urine from dogs treated with faropenem

The herein-described animal experiments were conducted under an ethics committee-approved protocol in accordance with the Tottori University Animal Use Committee (approval number: 15-T-46) and carried out as described previously [14]. Six beagle dogs (4 male, 2 female; mean age and weight, 6.3 ± 3.7 years and 12.4 ± 1.18 kg, respectively) were purchased from Kitayama Labes Co., Ltd. (Nagano, Japan). Prior to this study, all dogs were confirmed to be clinically healthy based on a physical examination, complete blood count, biochemical blood test and urinalysis. A balloon catheter was placed in the urinary bladder of each dog to allow urine collection. The dogs were orally administered faropenem (Farom Dry Syrup for Pediatric®; Maruho Co., Ltd., Osaka, Japan) at a dose of 5 mg/kg body weight. Whole urine was obtained via the catheter at 4, 8, and 12 h after administration. The samples were sterilized by filtration and stored at −80°C until analysis.

### Measurement of urine faropenem concentration with LC-MS

Reference standard faropenem and cephalexin as the internal standard were separately dissolved in acetonitrile and then diluted with ultrapure water. LC-MS was carried out with a high-performance liquid chromatograph (LC-10AT; Shimadzu Co., Ltd., Kyoto, Japan). The mass spectrums of faropenem and cephalexin were represented by peaks at *m/z* 308.0553–308.0558 and *m/z* 352.10, respectively. The compounds were separated on a 2.1-mm internal diameter × 100-mm length, 3-μm analytical column operated at 40°C (Mastro C18; Shimadzu GLC Ltd., Tokyo, Japan). The mobile phase comprised 0.1% formic acid aqueous solution and acetonitrile, and the flow rate was 0.2 mL/min. The injection volume was 0.1 μL. Standard samples for creation of a calibration curve were prepared with blank urine matrix spiked with six concentrations of faropenem (1, 5, 10, 50, 100, and 500 μg/mL). Standard and dog urine samples (50 μL) were mixed with 100 μg/mL of cephalexin (50 μL) as the internal standard and methanol (400 μL). After centrifugation at 12,000 rpm for 5 min, the supernatants were harvested and then diluted 10-fold with ultrapure water for analysis. The validity of the LS-MS assay was verified according to the guideline provided by the US Food and Drug Administration [15].

### Test organisms

The six ESBL-EC strains (strains ES-EC1–ES-EC6) from dogs with UTIs were selected from our collection [7] and used in this study. The faropenem minimum inhibitory concentrations (MICs) of these strains were determined in our previous study [7]. According to the tentative breakpoint set by Fuchs et al. [16], the six strains were categorized as follows: strains ES-EC1 to ES-EC4, susceptible; strain ES-EC5, intermediately susceptible; and strain ES-EC6, resistant (Table 1).

**Table 1.** MIC, MBC, and MUBC of faropenem for the six ESBL-producing *E. coli* strains tested in this study.

| Strain | MIC<sup>a</sup><br>(μg/mL) | MBC<br>(μg/mL) | MUBC (μg/mL) | | MUBC/MBC |
| --- | --- | --- | --- | --- | --- |
|  |  |  | Median | Range |  |
| ES-EC1 | 1 (S) | 2 | 1.29 | (0.64-3.57) | 0.64 |
| ES-EC2 | 1 (S) | 1 | 0.98 | (0.12-10.46) | 0.98 |
| ES-EC3 | 2 (S) | 4 | 4.68 | (1.0-14.7) | 1.17 |
| ES-EC4 | 2 (S) | 2 | 1.81 | (0.66-4.16) | 0.91 |
| ES-EC5 | 4 (I) | 4 | 14.5 | (3.1-20.9) | 3.62 |
| ES-EC6 | 16 (R) | 16 | 11.94 | (5.56-28.57) | 0.75 |

<sup>a</sup>S, susceptible; I, intermediate; and R, resistant according to Fuchs et al. [16].
MIC, minimum inhibitory concentration; MBC, minimum bactericidal concentration; MUBC, minimum urinary bactericidal concentration; ESBL, extended-spectrum β-lactamase

MIC, minimum inhibitory concentration; MBC, minimum bactericidal concentration; MUBC, minimum urinary bactericidal concentration; ESBL, extended-spectrum β-lactamase

### Determination of minimum bactericidal concentration

The minimum bactericidal concentration (MBC) was defined as the minimum concentration of drug needed to kill ≥99.9% of viable organisms after incubation for 24 h, according to the Clinical and Laboratory Standards Institute guideline [17].

### Determination of urinary bactericidal titer, area under the UBT–time curve, and minimum urinary bactericidal concentration

The urinary bactericidal titers (UBTs) corresponded to the maximal dilution titer of urine allowing bactericidal activity and were determined as described previously [14,18]. A logarithmic serial two-fold dilution was prepared using a 1:1 mixture of the urine sample obtained every 4 h after administration (see section titled “Sampling of urine from dogs treated with faropenem”) and the individual dog’s antimicrobial-free urine obtained prior to drug administration. UBTs were determined using a microdilution test system. Each well of the microplates contained 100 μL of the prepared dilution. The final inoculum was about 5 × 10^5^ CFU/mL. The plates were incubated at 35°C for 18 h, and the subcultured urine was then transferred to antimicrobial-free agar. The plates were incubated at 35°C overnight. The number of colonies subsequently grown was used to determine the bactericidal endpoint. The UBT was defined as a ≥99.9% reduction of the initially inoculated colony counts. A UBT of 0 was defined as no bactericidal activity, and a UBT of 1 was assigned when only undiluted urine displayed bactericidal activity. UBTs were transformed into ordinal data and described with reciprocal numbers [13,14].

The area under the UBT–time curve (AUBT) was calculated as the sum of the products of the reciprocal UBT values and the respective time (h) intervals for each test organism to easily compare UBT data among the tested strains. Calculation of the AUBT is an approximation considering the 4-h time intervals and the nonlinear kinetics in urine [13,14].

The minimum urinary bactericidal concentration (MUBC) for each strain was determined by dividing the antimicrobial concentration in a urine sample by the corresponding UBT [14,18].

### Statistical analysis

Repeated analysis of variance with Bonferroni correction was used to compare urine concentrations between collecting time periods. The median values of UBT, AUBT, and MUBC among the six dogs were calculated from the average value of the two middle elements. The UBTs for each strain were compared between collecting time periods by Friedman’s test followed by Scheffe’s test. The AUBTs were compared among the six tested strains by the Tukey–Kramer test. Spearman’s rank correlation coefficient (*ρ*) was calculated between the MIC and median AUBT. A *P*-value of <0.05 was considered significant for all analyses.

## Results

### Safety and laboratory test results

No adverse effects were observed in any dogs during the test period. The results of the physical examination, complete blood count, and biochemical blood test displayed no clinically relevant changes.

### Urine concentration and urinary excretion

The LC-MS assay showed a lower limit of quantitation at 10 ng/mL for faropenem in dog urine. The urine volume, urinary concentration, and cumulative excretion are shown in Table 2. The mean urinary concentration peaked 0 to 4 h after administration (584 µg/mL), then decreased to 246 µg/mL at 4 to 8 h and 23 µg/mL at 8 to 12 h. A significant difference in the urine concentration was observed between 0 to 4 h and 8 to 12 h (*P* < 0.05). All urine samples collected prior to drug administration had no detectable drug. The mean urinary excretion was 25.3% at 0 to 4 h and then remained almost unchanged from 4 to 8 h (34.3%) to 8 to 12 h (35.5%).

**Table 2.** Urinary volume, concentration, and cumulative excretion of faropenem in six dogs.

| Collection period (h) | Vol (ml) | Conc (µg/mL) | Cumulative urinary excretion (%) |
| --- | --- | --- | --- |
| 0-4 | 44 (13-78) | 584 (147-1748) | 25.3 (11.9-48.3) |
| 4-8 | 28 (13-50) | 246 (64-669) | 34.3 (19.1-66.9) |
| 8-12 | 32 (13-67) | 23 (LLQ-35) | 35.5 (19.1-67.3) |

Data are presented as mean (range).

Data are presented as mean (range).

Vol, volume; Conc, concentration; LLQ, lower level of quantification.

### UBTs and AUBTs

The temporal changes in the median UBTs for each strain are shown in Figure 1. In all tested ESBL-EC strains, the median UBTs of faropenem peaked at 0 to 4 h and then significantly decreased at 4 to 8 h and/or 8 to 12 h (*P* < 0.05).

**Fig. 1.**
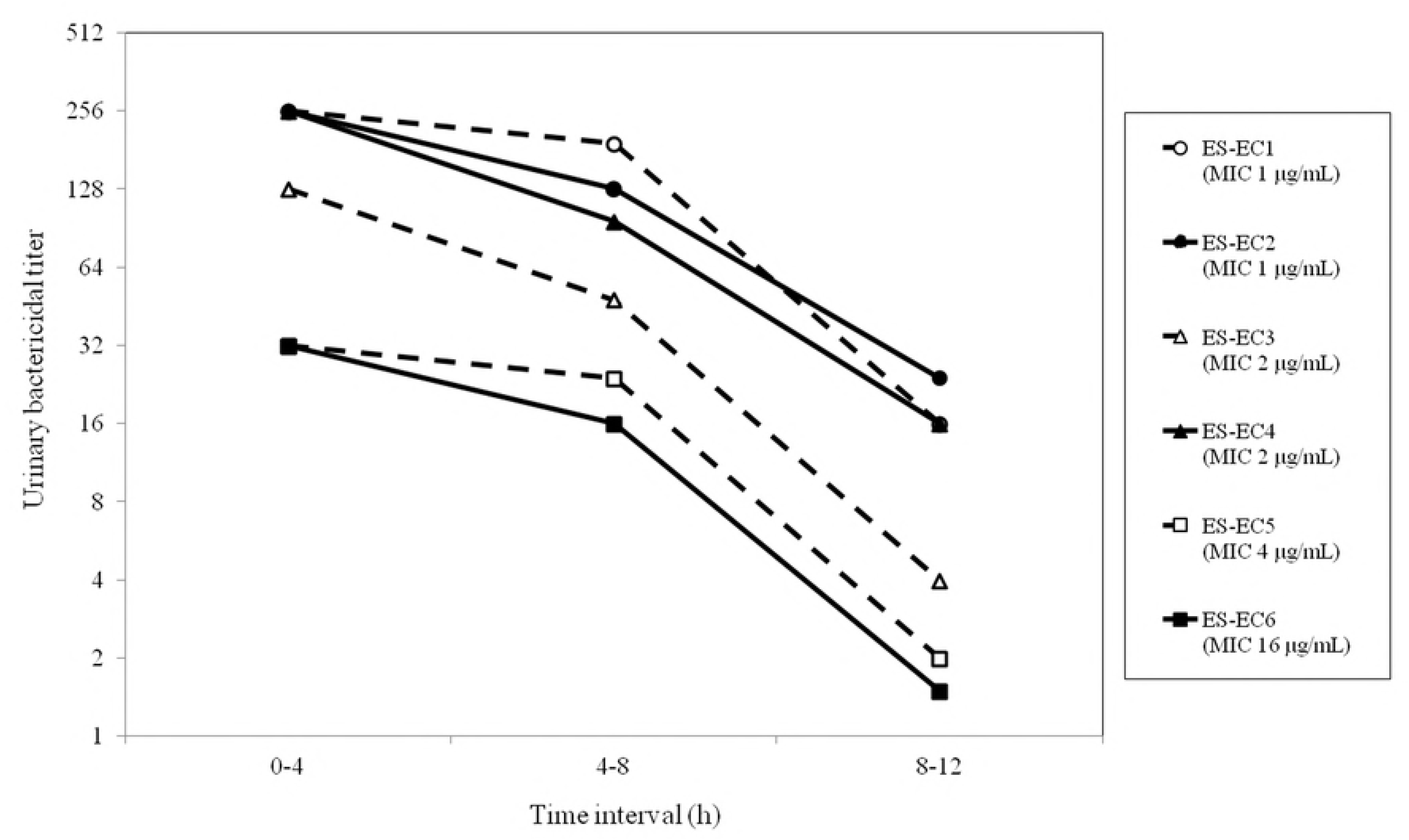
Reciprocal UBTs of faropenem (5 mg/kg body weight) for the six ESBL-producing *Escherichia coli* strains tested in this study. UBT, urinary bactericidal titer; ESBL, extended-spectrum β-lactamase; MIC, minimum inhibitory concentration.

Of the tested strains, the two strains with an MIC of 1 µg/mL (ES-EC1 and ES-EC2) had significantly higher median AUBTs (1072–1312) than the strains with an MIC of 4 µg/mL (ESEC5) and 16 µg/mL (ES-EC6) (*P* < 0.05) (Fig. 2).

**Fig. 2.**
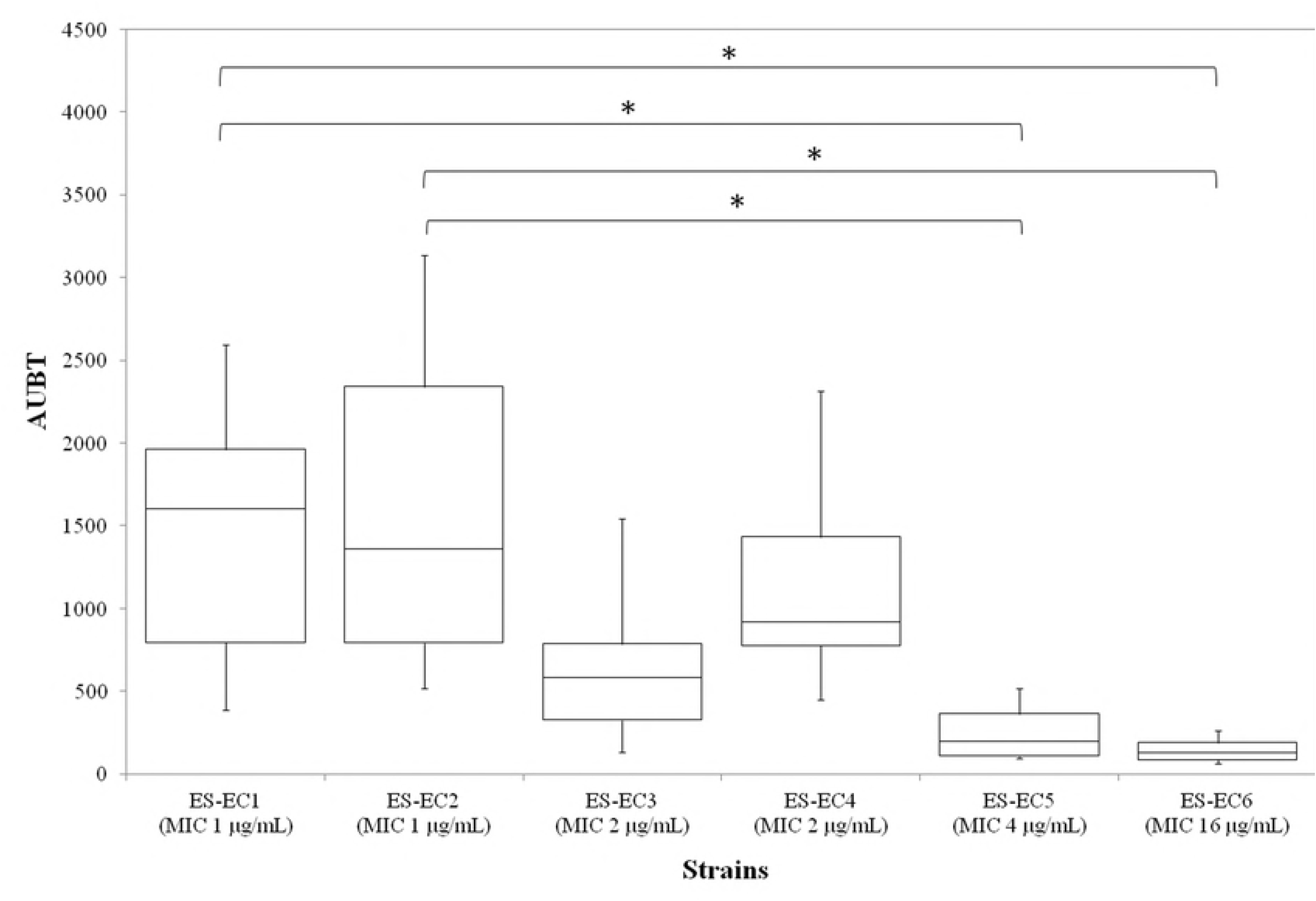
Comparison of AUBTs of faropenem between the six ESBL-producing *Escherichia coli* strains tested in this study. AUBT, area under the urinary bactericidal titer–time curve; ESBL, extended-spectrum β-lactamase; MIC, minimum inhibitory concentration. *Significant differences between groups (*P* < 0.05).

Spearman’s rank correlation coefficient between the MICs and median AUBTs was −0.968 (*P* < 0.05).

### MBCs and MUBCs

The MBCs ranged from 1 to 16 µg/mL (one to two times the corresponding MIC for each strain), whereas the median MUBCs ranged from 0.98 to 11.94 µg/mL (Table 1). The ratios of the median MUBC to the corresponding MBC ranged from 0.64 to 3.62.

## Discussion

Although faropenem shows excellent *in vitro* antimicrobial activity against ESBL-producing bacteria [7], its efficacy for canine UTIs with these bacteria has not been previously assessed. To our knowledge, this is the first report to investigate the urinary pharmacokinetics and pharmacodynamics of faropenem in dogs.

The present study demonstrated that in dogs, approximately one-third of the dose of faropenem is excreted in urine 12 h after oral administration. This urinary excretion of faropenem in dogs is higher than that in humans: urinary excretion of the drug after oral administration at a 300-mg dose in human patients was merely 14% to 20% [9]. These findings imply that this drug is more suitable for treatment of UTIs in dogs than in humans, possibly because of the differences in absorption, distribution, metabolism, and elimination between dogs and humans.

In this study, we also found an extremely low concentration of faropenem in dog urine at 8 to 12 h, and the cumulative excretion remained almost unchanged from 4 to 8 h to 8 to 12 h. This indicates that urinary excretion of faropenem practically expires at 12 h after oral administration in dogs. This urinary pharmacokinetic property of faropenem differs greatly from that of once-a-day fluoroquinolones, which can maintain high urinary concentrations until 24 h after oral administration [14,19,20]. In addition, the UBTs, which can serve as a pharmacokinetic/pharmacodynamic assessment parameter of antimicrobial agents in the urine [13,14], of faropenem in all tested strains fluctuated closely with the urine concentration of the drug during the same period; notably, the UBTs significantly decreased 8 to 12 h after oral administration. Our results may indicate that administration of faropenem two or three times a day is more appropriate than once a day for the treatment of canine UTIs.

When comparing UBTs among the six ESBL-producing *E. coli* strains tested in this study, the strains with the lower MICs generally exhibited the higher UBTs of faropenem during the test periods. A similar finding was confirmed in our previous study on the UBTs of orbifloxacin [14]. Additionally, the AUBTs, which reflect the overall UBTs, were greatly dependent upon the respective MIC of each strain. Notably, the AUBTs of the two strains with a faropenem MIC of 1 µg/mL, which is the minimum concentration required to inhibit 90% of ESBL-producing *E. coli* [7], were extremely high. This implies that most ESBL-EC–related UTIs in dogs can be theoretically treated with faropenem at an oral dose of 5 mg/kg body weight. Compared with the two susceptible strains with an MIC of 1 µg/mL, the strains with an MIC of 4 and 16 µg/mL had significantly lower AUBTs (approximately 1/10 of those of the susceptible strains), suggesting that treatment with faropenem at the same dose has less therapeutic efficacy for UTIs by *E. coli* strains with an MIC of ≥4 µg/mL. Fuchs et al. [16] proposed that the susceptibility breakpoint of faropenem is ≤2 µg/mL based on human pharmacokinetic parameters. The present findings suggest that this breakpoint is also reasonable for canine UTIs when treated with faropenem at a dose of 5 mg/kg. Further clinical studies are needed to establish this value as the valid clinical breakpoint.

Like the study by Matsuzaki et al. [21], our data showed a low MBC/MIC ratio (within one dilution), indicating that faropenem has *in vitro* bactericidal activity against ESBL-EC. In the present study, we calculated the MUBCs in each test strain based on the urine concentration and UBTs to assess the activity of faropenem in dog urine. As a result, the median MUBCs were 0.6- to 3.6-fold higher than the corresponding MBCs. This MUBC/MBC ratio was relatively low compared with orbifloxacin, for which the MUBC/MBC ranged from approximately 2 to 15 [14]. Comparatively, therefore, the antimicrobial activity of faropenem might be minimally decreased in dog urine.

## Conclusion

We determined the UBTs and related parameters of faropenem in dogs to assess the efficacy of this drug against canine UTIs with ESBL-producing bacteria. Based on the urinary pharmacokinetics and UBTs of faropenem, administration more than once daily is recommended for treatment of these UTIs in dogs. In addition, the comparison of AUBTs among the strains with different MICs might suggest that an MIC of ≤2 µg/mL is applicable to the susceptible breakpoint of faropenem for canine UTIs when administered at 5 mg/kg body weight. We strongly believe that the present study serves as a basis for clinical application of faropenem for ESBL-producing bacteria-related UTIs in dogs.

## Competing interests

K.H. received JPSP KAKENHI Grant Number 16K18804 (Japan Society for the Promotion of Science; https://www.jsps.go.jp/j-grantsinaid/). The funder had no role in the study design, data collection and analysis, decision to publish, or preparation of the manuscript.

## Acknowledgments

We thank Ms. Mizuki Yokono for providing technical advice concerning LC-MS and Dr. Taku Tsukamoto for providing the analytical column for LC-MS. We also thank Angela Morben, DVM, ELS, from Edanz Group (www.edanzediting.com/ac), for editing a draft of this manuscript.

